# Distinct prophase arrest mechanisms in human male meiosis

**DOI:** 10.1101/195321

**Authors:** Sabrina Z. Jan, Aldo Jongejan, Cindy M. Korver, Saskia K. M. van Daalen, Ans M. M. van Pelt, Sjoerd Repping, Geert Hamer

## Abstract

To prevent chromosomal aberrations to be transmitted to the offspring, strict meiotic checkpoints are in place to remove aberrant spermatocytes. However, in about 1% of all males these checkpoints cause complete meiotic arrest leading to azoospermia and subsequent infertility. We here unravel two clearly distinct meiotic arrest mechanisms that act during the prophase of human male meiosis. Type I arrested spermatocytes display severe asynapsis of the homologous chromosomes, disturbed XY-body formation and increased expression of the Y-chromosome encoded gene *ZFY* and seem to activate a DNA damage pathway leading to induction of p63 mediated spermatocyte elimination. Type II arrested spermatocytes display normal chromosome synapsis, normal XY-body morphology and meiotic crossover formation but have a lowered expression of several cell cycle regulating genes and fail to properly silence the X-chromosome encoded gene *ZFX*. Discovery and understanding of these meiotic arrest mechanisms increases our knowledge on how genomic stability is guarded during human germ cell development.

---

While our somatic bodies inevitably die of old age or disease, our germ cells have to maintain sufficient genome integrity to pass on our genome to, in principal, endless generations. Therefore, to prevent transmission of aneuploidies or other chromosomal aberrations, strict genome integrity checkpoints exist in the process of meiosis to remove germ cells that fail certain quality checks.

During meiosis, in order to generate haploid sperm or oocytes, diploid germ cells undergo two consecutive rounds of chromosome segregation after one round of DNA replication. During meiosis I the homologous chromosomes, each consisting of one pair of sister chromatids, are segregated, followed by separation of the sister chromatids into haploid cells during meiosis II. Successful meiosis requires that the homologous chromosomes are properly paired and aligned. This is achieved by the induction of DNA double-strand breaks (DSBs) during the prophase of the first meiotic division by the protein SPO11. The repair of these SPO11-induced DSBs initiates and requires synapsis of the homologous chromosomes and ensures the formation of at least one meiotic crossover per homologous chromosome pair ^1^.

In the mouse, failure to properly repair meiotic DSBs or synapse the homologous chromosomes leads to arrest during the first meiotic prophase at a specific stage of spermatogenesis, the so called epithelial stage IV arrest ^2, 3^. However, despite displaying spermatocyte apoptosis at the same stage of spermatogenesis, different meiotic recombination mouse mutants show different responses and cytological end-points ^4^. This led to the idea that more than one checkpoint mechanism exists that can induce apoptosis of meiotic cells at stage IV of mouse spermatogenesis.

One type of mouse stage IV arrest occurs independent of SPO11-induced DSBs ^5, 6^ or the conventional DNA damage response protein p53 ^7, 8^, and is caused by incomplete synapsis of the homologous chromosomes ^9, 10^. When homologous chromosomes synapse, the checkpoint protein TRIP13 removes the meiosis specific HORMA-domain proteins HORMAD1 and HORMAD2 from the chromosome axes ^11^. However, on asynapsed chromosome axes these proteins remain present and recruit the kinase ATR ^12, 13^, which together with several other proteins such as BRCA1 and γH2AX, mark the silencing of transcription from asynapsed chromosomal regions via a process referred to as meiotic silencing ^14, 15^. Usually, when the autosomes are fully synapsed, only the X and Y-chromosomes are subject to meiotic silencing because they remain largely unsynapsed due to a lack of sequence homology. This leads to the formation of the XY-body in which the sex chromosomes are bound to i.e. ATR, BRCA1 and γH2AX ^9, 10^. However, in case of extensive autosomal asynapsis, these proteins are sequestered away from the sex chromosomes. This, in turn, leads to failure to silence the Y-chromosomal genes *Zfy1* and *Zfy2* that, via a yet unknown mechanism, induce apoptosis of spermatocytes at stage IV of spermatogenesis ^16^.

Because synapsis of the homologous chromosomes and meiotic recombination are two highly intertwined events, i.e. problems with recombinational repair will most often also lead to asynapsis and vice versa, the possibility that two separate meiotic checkpoints may act at stage IV of spermatogenesis has long been overlooked. However, it has been found that the canonical DNA damage response pathway, consisting of i.e. MRE11, NBS1, ATM and the checkpoint kinase CHK2, can induce mouse spermatocyte apoptosis prior to XY-body failure induced apoptosis ^17^. Moreover, in mouse oocytes, in which apoptosis cannot be induced via *Zfy*-expression due to the absence of a Y-chromosome, unrepaired meiotic DSBs also activate CHK2 which, subsequently, provokes apoptosis via the DNA damage response proteins p53 and p63 ^18-20^. Indeed, both p53 and p63 are also present in mouse spermatocytes ^21, 22^ and have been recently found to be specifically involved in recombination-dependent pachytene arrest of mouse spermatocytes ^23^.

In contrast to the mouse, the mechanisms of human meiotic arrest have not been thoroughly investigated at the molecular level and are poorly understood. Nevertheless, about 10-20% of men with non-obstructive azoospermia are diagnosed with, complete or incomplete, meiotic arrest ^24, 25^. Some of these men are carriers of a known genetic aberration, for instance a defined chromosomal translocation or duplication ^26^ or a single gene mutation that has been associated with meiotic arrest ^27-31^, but the etiology remains unknown in the vast majority of men. Given the hundreds of genes or unknown environmental factors that may be involved, and the lack of appropriate human genetic models, a general meiotic arrest mechanism has not been determined in humans.

## Results

### Two types of human meiotic prophase arrest

To investigate human meiotic arrest we collected testis biopsies from men with non-obstructive azoospermia diagnosed with maturation arrest. From 2011 till 2013, we collected 350 testicular biopsies, of which 14 displayed complete meiotic arrest. After careful histological examination, four samples were deemed unfit for single cell laser dissection microscopy (LDM) due to poor morphology of the testicular sections. The remaining 10 patients were used for this study. In these patients we first investigated whether the arrested spermatocytes had formed a normal XY-body. We therefore stained paraffin embedded testis sections with an antibody against γH2AX (Fig. 1). Before completion of synapsis of the homologous chromosomes at the zygotene stage, γH2AX marks all asynapsed chromosome axes. Subsequently, in healthy pachytene spermatocytes, γH2AX becomes restricted to the XY-body in which the X and Y chromosomes are not fully synapsed and transcriptionally silenced ^9, 10, 15^. In five patients, hereafter referred to as type I, meiotic prophase arrest was characterized by the absence of a discernable XY-body and γH2AX staining dispersed throughout the nucleus. The other five men, hereafter referred to as type II, also displayed meiotic prophase arrest but showed similar γH2AX staining as in controls with normal spermatogenesis (Fig. 1).

**Figure 1.**
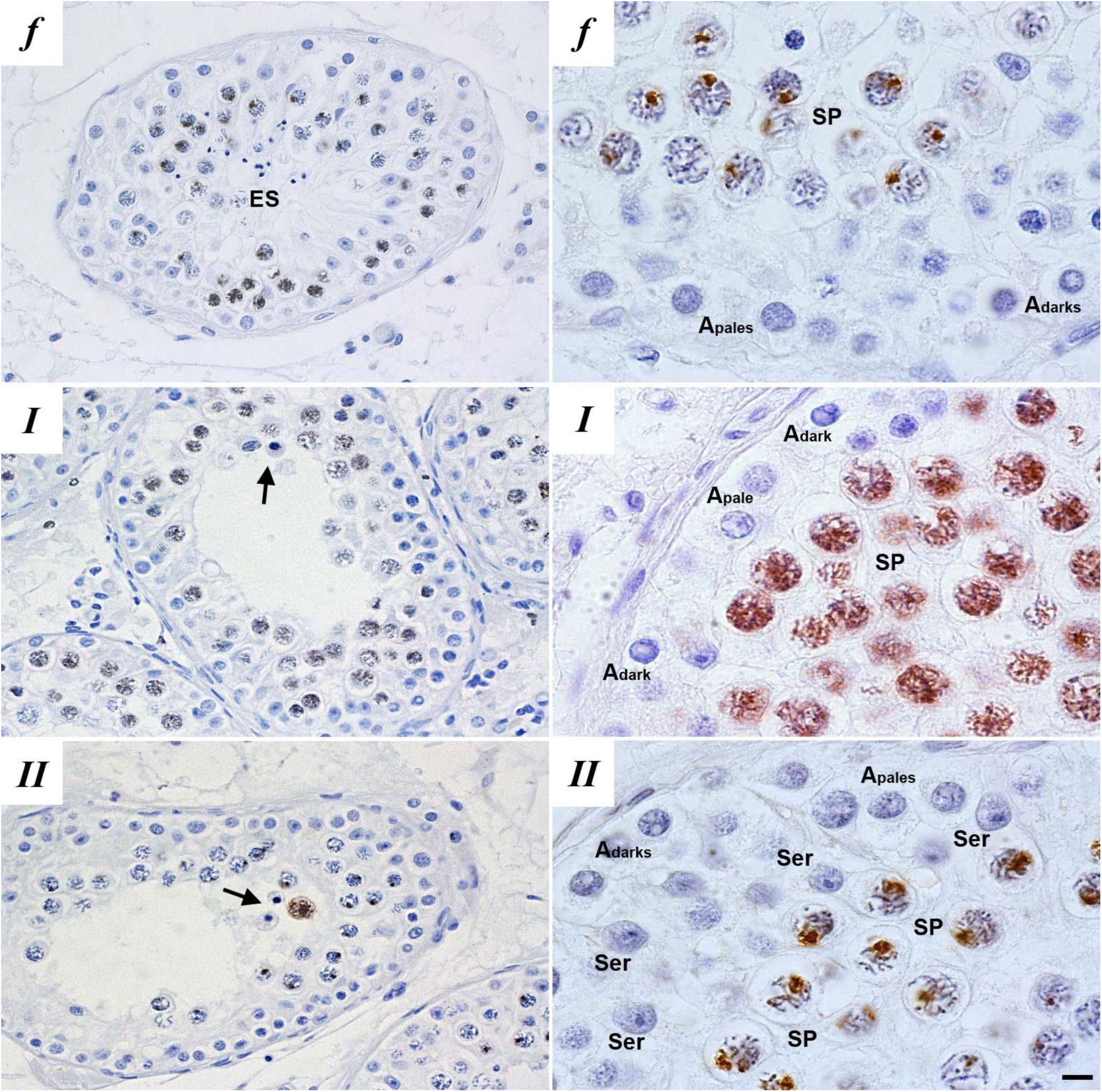
Histological evaluation and immunohistochemical localization of γH2AX in paraffin embedded testis sections of fertile men (***f***) and patients with meiotic maturation arrest reveal two types of meiotic prophase arrest patients: type I (***I***) and type II (***II***) meiotic arrest. Type I (***I***) meiotic arrest patients display meiotic prophase arrest and disturbed γH2AX distribution and no XY-body formation, type II (***II***) meiotic arrest patients display meiotic prophase arrest but normal γH2AX distribution and XY-body formation. Depicted are: Sertoli cells (Ser), elongated spermatids (ES), A_pale_ and A_dark_ spermatogonia, spermatocytes (SP) and apoptotic spermatocytes (arrows). Bar = 10 μm.

We then made meiotic spread preparations from biopsies of the same patients to stain for γH2AX and the synaptonemal complex protein SYCP3 to further investigate XY-body formation and homologous chromosome synapsis. In contrast to controls with normal spermatogenesis, type I spermatocytes displayed severe asynapsis of the homologous chromosomes, characterized by a zygotene-like appearance of SYCP3, and absence of a XY-body, marked by dispersed γH2AX staining covering all asynapsed chromosomes (Fig. 2a, *I*). Type II spermatocytes, on the other hand, reached full chromosome synapsis and formed XY-bodies similar to controls with normal spermatogenesis (Fig. 2a, *II*). In the mouse γH2AX has been shown to mark meiotic silencing of asynapsed chromosomes ^9, 10, 15^. To investigate whether this is also the case in human spermatocytes we combined γH2AX with an RNA staining protocol for Cot-1 to mark RNA-synthesis. Using confocal microscopy and max projection of the confocal layers, we found that, also in human spermatocytes, transcriptionally silent regions are marked by γH2AX (Extended Data Fig. 1). Moreover, we confirmed that type I spermatocytes did not form γH2AX positive XY-bodies normally marking the transcriptionally silent sex chromosomes.

**Figure 2.**
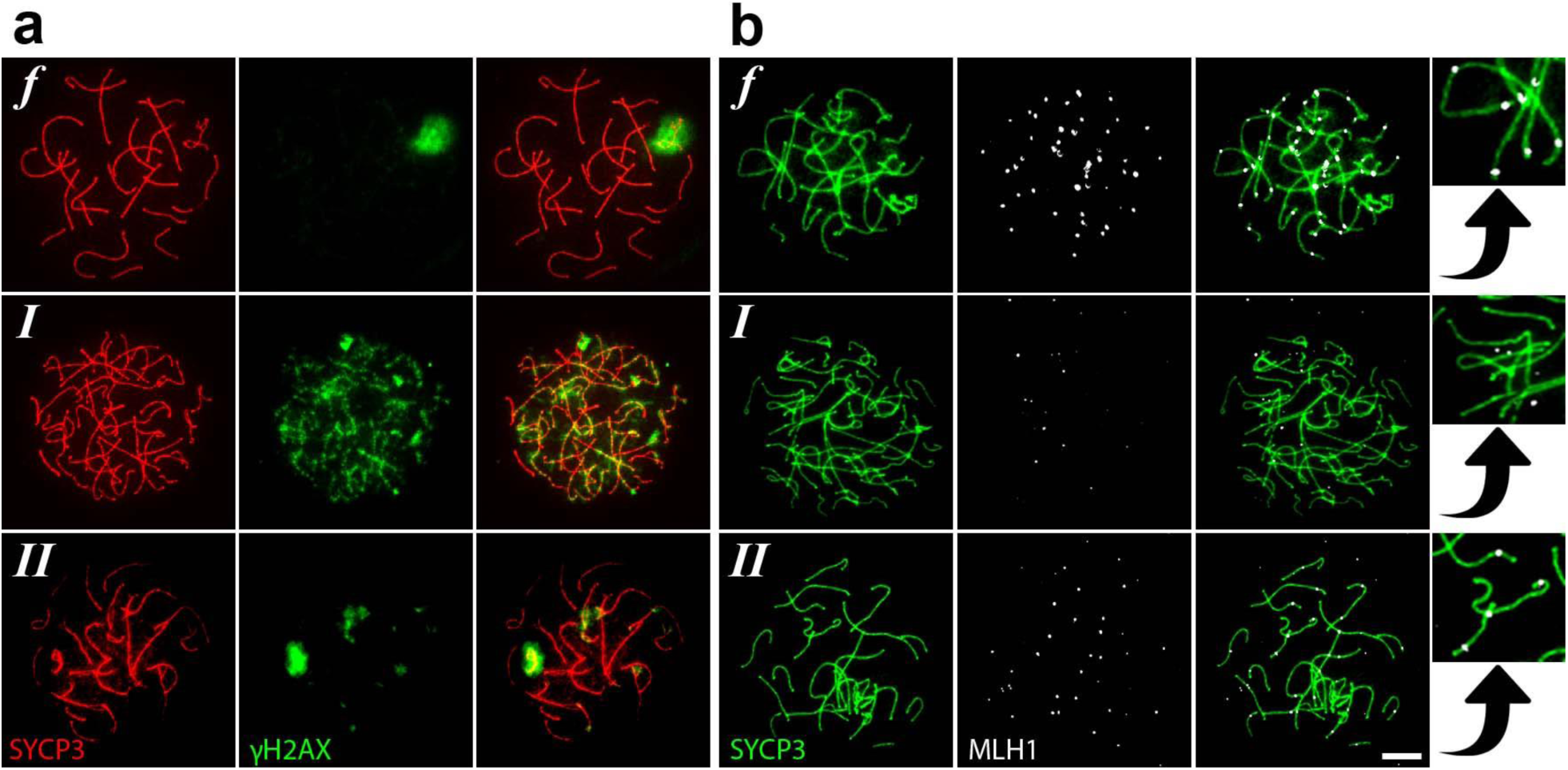
Immunofluorescent stainings of meiotic spread preparations of fertile men (***f***) and type I (***I***) and type II (***II***) meiotic arrest patients reveal severe asynapsis (SYCP3) (**a, b**), disturbed XY-body formation (γH2AX) (**a**) and absence of meiotic crossovers (MLH1) (**b**) in specifically type I meiotic arrest patients. Bar = 5 μm.

To further evaluate meiotic progression, we used staining against MLH1 to investigate whether meiotic crossover formation is disturbed. As expected type I patients showed zygotene-like spermatocytes with asynapsed homologous chromosome and no MLH1 staining on the chromosome axes (Fig. 2b, *I*). However, like in the controls with normal spermatogenesis, all type II patients showed pachytene spermatocytes with fully synapsed homologous chromosomes and MLH1 foci marking meiotic crossover sites (Fig. 2b, *II*).

Thus, based on these histological and cytological evaluations of testis samples of men with meiotic arrest from our clinic we identified the existence of two types of human meiotic prophase arrest. One group shows aberrant XY-body formation, severe asynapsis of the homologous chromosomes and meiotic arrest comparable to stage IV arrest in mouse. A second group displays normal chromosome synapsis, XY-body morphology and crossover formation and is distinct from stage IV meiotic arrest in mouse.

### Different types of arrested spermatocytes have distinct gene expression profiles

In order to try to understand the molecular mechanisms underlying these different types of meiotic arrest, we used a protocol for single cell LDM and RNA-sequencing ^32^ to generate the transcriptomic profiles of pachytene spermatocytes from the ten patients with meiotic arrest. Five-hundred morphologically normal and non-apoptotic pachytene spermatocytes per patient were isolated and pooled for further RNA-sequencing and analysis. Comparison of these profiles with expression profiles of pachytene spermatocytes from a similar spermatogenic stage but derived from men with normal spermatogenesis ^32^, revealed that arrested pachytene spermatocytes are transcriptomically distinct from normal pachytene spermatocytes. At the transcriptome level, most arrested spermatocytes appeared to be more leptotene/zygotene-like than pachytene-like (Fig. 3a). Notably, type I spermatocytes clustered more closely together, whereas we observed more biological variation between type II spermatocytes (Fig. 3a). Spermatocytes derived from one type II patient clustered closely to controls with normal spermatogenesis (Fig. 3a). We then evaluated more testis sections of all patients and found that that spermatogenesis of specifically this patient progresses beyond the first meiotic prophase and arrests later at a meiotic metaphase stage (Extended Data Fig. 2). Because only a single patient displayed this type (III) of arrest we decided to exclude this patient from further downstream analysis to avoid drawing too far reaching conclusions based on a single case.

**Figure 3.**
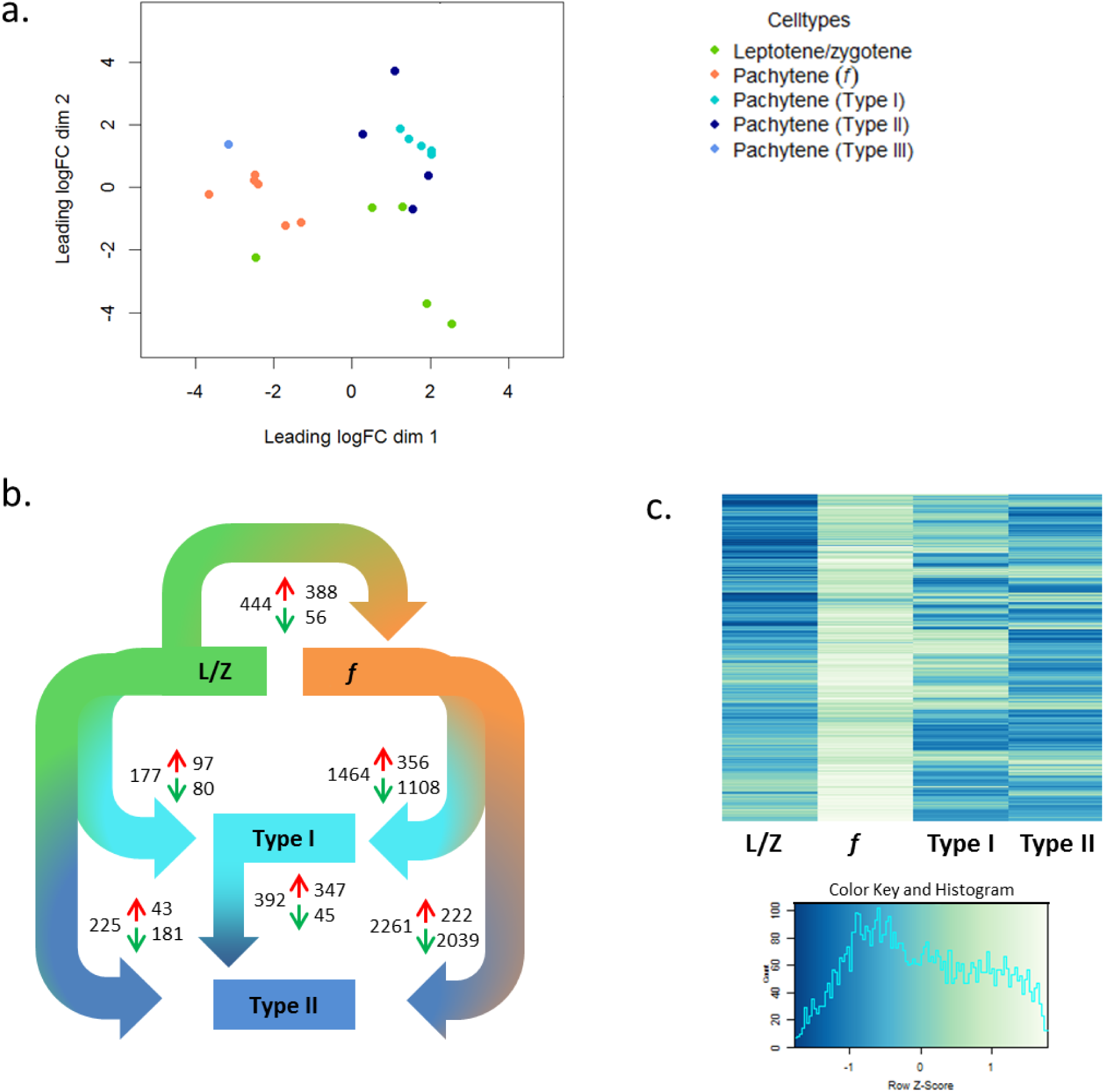
Transcriptomic analysis of fertile, type I and II spermatocytes. Multidimensional scaling of leptotene/zygotene spermatocytes (L/Z, green) and pachytene spermatocytes from fertile men (*f*, coral) alongside type I (Type I, turquoise), type II (Type II, dark blue) and type III (Type III, light blue) arrested pachytene spermatocytes (**a**) and differential gene expression analysis (adjusted p-value < 0,05; **b**) reveal distinct transcriptomic profiles for type I and II arrested spermatocytes compared to leptotene/zygotene (L/Z) and pachytene spermatocytes (*f*) from fertile men. Red arrow; genes upregulated, green arrow; genes downregulated. (**c**) Analysis of genes upregulated during the leptotene/zygotene to pachytene transition in normal spermatogenesis reveals a leptotene/zygotene-like expression pattern in type I and II arrested spermatocytes.

Differential gene expression (DEG) analysis confirmed that both type I and II arrested spermatocytes more closely resemble leptotene/zygotene-like cells than normal pachytene spermatocytes (Fig. 3b). We therefore looked more closely at the expression of genes that we previously found to be upregulated during the leptotene/zygotene to pachytene transition ^32^. This analysis first revealed that many genes from this gene set were not upregulated in arrested pachytene spermatocytes but remained at leptotene/zygotene-like expression levels (Fig. 3c). Moreover, the two types of meiotic prophase arrest again appeared clearly distinct, showing different sets of genes that fail to reach normal pachytene expression levels (Fig. 3c).

### The two types of meiotic prophase arrest are characterized by specific biological processes

Subsequent k-means clustering of the DEGs, based on their expression profile in fertile, type I or type II spermatocytes, generated eight major gene expression clusters (Fig. 4, Supplementary table 1). To be able to investigate the differences in molecular pathways between the two patient groups we performed a gene ontology analysis using DAVID on these clusters (Table 1, **Supplementary table 2**).

**Table 1:** Gene set enrichment analysis (GO-terms) of DEGs in each k-means cluster

| Cluster | Representative GO term for enriched gene sets | Enrichment Score |
| --- | --- | --- |
| **1** | GO:0006325~chromatin organization | 11.24 |
|  | GO:0016570~histone modification | 2.95 |
|  | GO:0010556~regulation of macromolecule biosynthetic process | 2.66 |
|  | GO:0016050~vesicle organization | 1.96 |
|  | GO:0006355~regulation of transcription, DNA-dependent | 1.83 |
|  | GO:0016570~histone modification | 1.63 |
| **2** | GO:0006810~transport | 1.72 |
|  | GO:0016192~vesicle-mediated transport | 1.70 |
| **3** | GO:0022402~cell cycle process | 2.96 |
|  | GO:0030163~protein catabolic process | 2.25 |
|  | GO:0006396~RNA processing | 1.68 |
|  | GO:0034623~cellular macromolecular complex disassembly | 1.61 |
|  | GO:0031396~regulation of protein ubiquitination | 1.48 |
|  | GO:0032886~regulation of microtubule-based process | 1.36 |
|  | GO:0030071~regulation of metaphase/anaphase transition | 1.32 |
|  | GO:0006461~protein complex assembly | 1.30 |
| **4** | no significant gene enrichment |  |
| **5** | GO:0030154~cell differentiation | 1.64 |
|  | GO:0060348~bone development | 1.53 |
| **6** | GO:0007283~spermatogenesis | 7.38 |
|  | GO:0016337~cell-cell adhesion | 5.98 |
|  | GO:0016339~calcium-dependent cell-cell adhesion | 2.87 |
|  | GO:0030317~sperm motility | 2.55 |
|  | GO:0006811~ion transport | 1.30 |
| **7** | GO:0045333~cellular respiration | 1.44 |
|  | GO:0007283~spermatogenesis | 1.38 |
| **8** | GO:0043170~macromolecule metabolic process | 4.40 |
|  | GO:0051276~chromosome organization | 4.14 |
|  | GO:0006396~RNA processing | 3.15 |
|  | GO:0010608~posttranscriptional regulation of gene expression | 2.19 |
|  | GO:0050658~RNA transport | 2.16 |
|  | GO:0008219~cell death | 2.14 |
|  | GO:0022414~reproductive process | 2.09 |
|  | GO:0044267~cellular protein metabolic process | 1.98 |
|  | GO:0010468~regulation of gene expression | 1.49 |
|  | GO:0006508~proteolysis | 1.46 |
|  | GO:0006974~response to DNA damage stimulus | 1.43 |
|  | GO:0033036~macromolecule localization | 1.42 |

**Figure 4.**
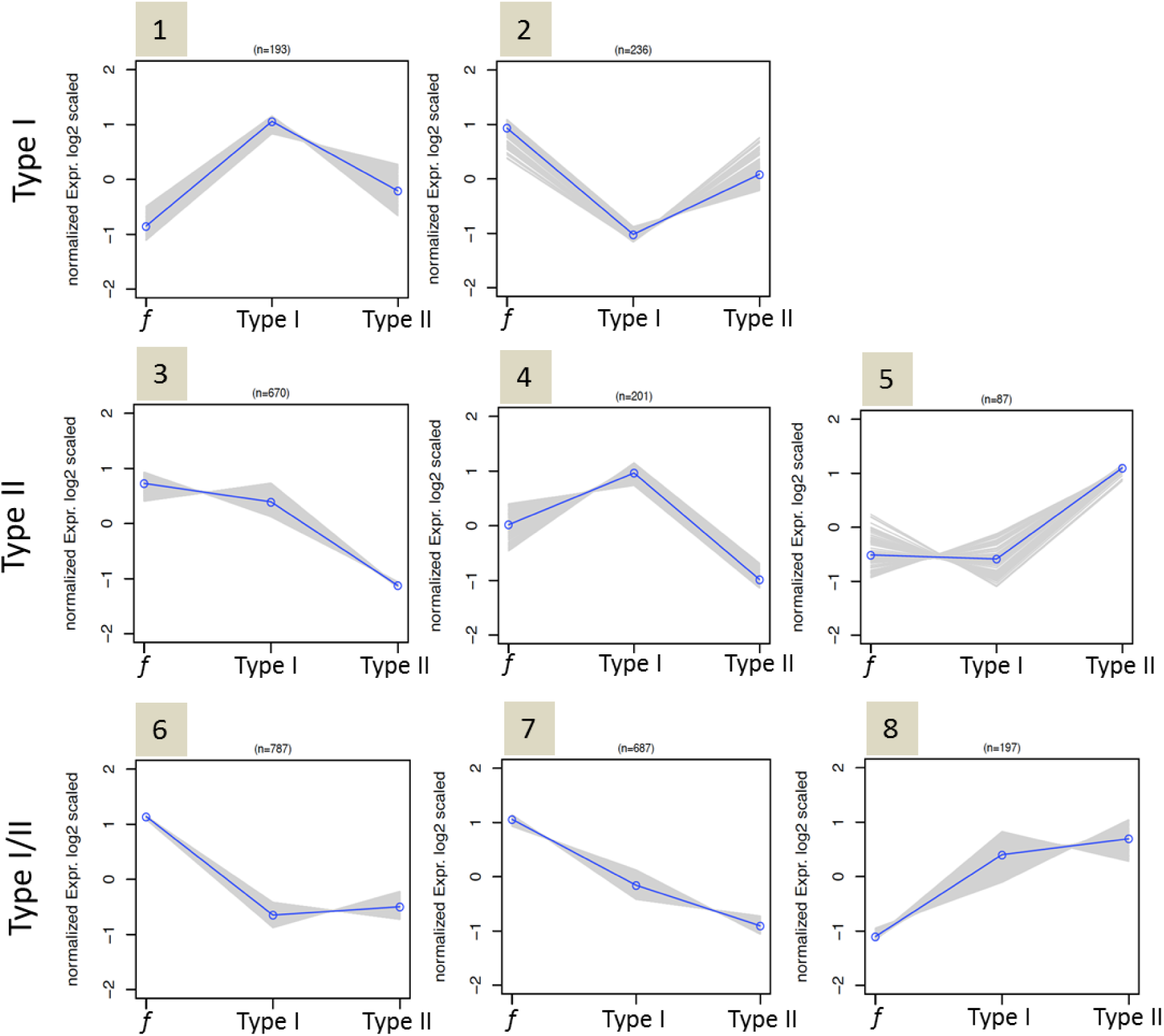
K-means cluster analysis of differentially expressed genes between fertile (*f*), type I and II spermatocytes (type I/II) reveals genes that are aberrantly expressed in type I spermatocytes (clusters 1, 2), type II spermatocytes (clusters 3, 4, 5) and in both type I and II spermatocytes (clusters 6, 7, 8).

Four-hundred-and-twenty-nine genes (clusters 1 and 2) were aberrantly expressed specifically in type I spermatocytes of which 193 genes were specifically upregulated in type I spermatocytes (cluster 1). Reflecting the clear defect in chromosome synapsis in type I spermatocytes, this cluster was highly enriched (enrichment score: 11.24 with >1.3 being significant) with a gene set representing chromatin organization. In addition, genes involved in histone modification were also clearly upregulated, as were genes involved in nucleic acid metabolic processes and gene transcription. The remaining 236 genes that were predominately downregulated in type I arrested spermatocytes (cluster 2) were enriched in genes involved in vesicle formation and transport.

Nine-hundred-and-fifty-eight genes (clusters 3, 4 and 5) were aberrantly expressed specifically in type II arrested spermatocytes, of which 871 genes were downregulated (clusters 3 and 4). This gene set included many genes involved in cell cycle progression, including the cyclins A1, A2 and E1, as well as genes involved in microtubule organization and the metaphase to anaphase transition. Also genes involved in macromolecule (protein) degradation and RNA processing were clearly over-represented in these clusters. The remaining 87 genes that were specifically upregulated in type II spermatocytes (cluster 5) were enriched for a gene set involved in cellular differentiation.

One-thousand-six-hundred-and-seventy-one genes were aberrantly expressed in both type I and II arrested spermatocytes (clusters 6, 7 and 8). 1474 genes were down regulated in both types of arrested spermatocytes (clusters 6 and 7). These genes appeared predominately involved in spermatogenesis, cell-cell adhesion and sperm motility. One-hundred-and-ninety-seven genes upregulated in both type I and type II spermatocytes (cluster 8) are involved in numerous processes, including macromolecule metabolic processes (for instance nucleic acid metabolic processes), chromosome structure and organization, RNA processing/transport, cell death and post-transcriptional regulation of gene expression.

In summary, both types of meiotic arrest displayed upregulation of genes involved in chromatin structure and organization and RNA processing. However, in type I spermatocytes chromatin structure and organization appeared to be the major underlying problem, while specifically type II meiotic arrest were characterized by downregulation of genes involved in RNA processing and cell cycle progression.

### Type I and II specific upregulation of the sex-chromosomes encoded genes *ZFY* and *ZFX*

Like in the mouse ^9, 10, 16^, type I meiotic prophase arrest could be caused by incomplete synapsis of the homologous chromosomes and subsequent failure to silence the sex chromosomes, leading to the expression of *Zfy*-genes and spermatocyte apoptosis. Also human type I spermatocytes show severe asynapsis of the homologous chromosomes and absence of a XY-body (Fig. 2) in which transcription is normally silenced (Extended Data Fig. 1). We therefore made beeswarm plots to visualize differential expression of the human *ZFY* gene in the spermatocytes of fertile men and men with type I and II spermatocyte arrest. Indeed, we found *ZFY* to be clearly upregulated in type I arrested spermatocytes (Fig. 5a). Hence, also during human male meiosis, asynapsis of the homologous chromosomes may lead expression of *ZFY* and subsequent spermatocyte elimination. Interestingly, the X-chromosome encoded gene *ZFX*, in the mouse expressed during the interphase between meiosis I and meiosis II ^33^, is specifically over-expressed in type II arrested spermatocytes (Fig. 5a) and may be thus be involved in the elimination of these cells.

**Figure 5.**
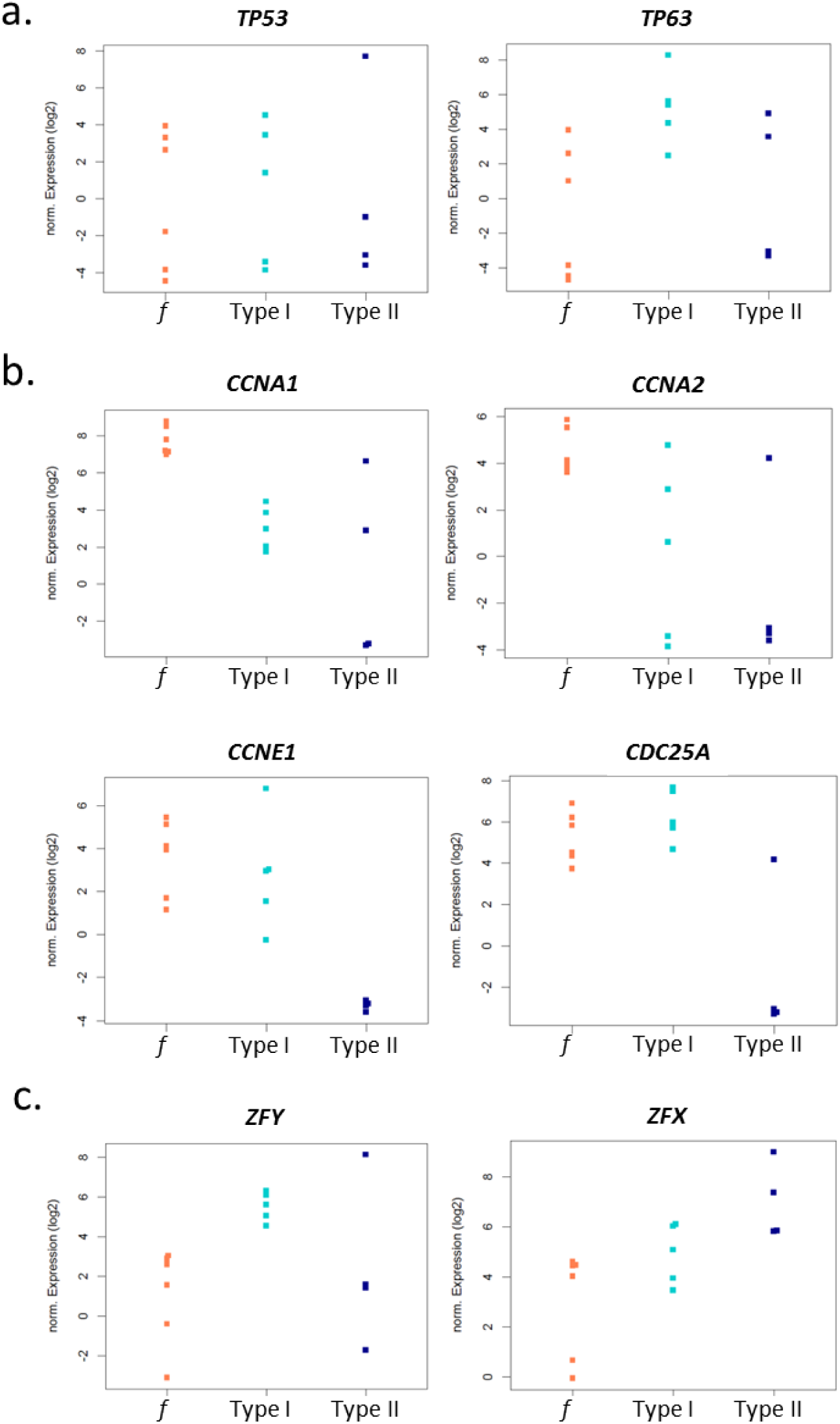
Beeswarms depicting different expression levels of *TP53* (adjusted p ≤ 0.8030) and *TP63* (adjusted p ≤ 0.0469) (**a**), *CCNA1* (adjusted p ≤ 0.0051), *CCNA2* (adjusted p ≤ 0.0626), *CCNE1* (adjusted p ≤ 0.0001) and *CDC25A* (adjusted p ≤ 0.0031) (**b**) and *ZFY* (adjusted p ≤ 0.0389)and *ZFX* (adjusted p ≤ 0.0899) (**c**) in fertile (*f*; coral), type I (type I; turquoise), type II (type II; dark blue) spermatocytes.

### Type I human arrested spermatocytes display upregulation of the DNA damage response protein p63

Alternatively type I meiotic arrest could be caused by the DNA damage pathway including ATM,CHK2 and the DNA damage response proteins p53 or p63 ^17-20^. We therefore made additional beeswarm plots to visualize differential expression of the human *TP53* and *TP63* genes (coding for p53 and p63) in fertile men and type I and II patients. The gene *TP53* did not detectably change in type I or II spermatocytes. However, we did find a strong upregulation of *TP63* in specifically type I spermatocytes (Fig. 5b). Hence, DNA damage checkpoint induced apoptosis of human type I spermatocytes is most likely mediated by activation of the DNA damage response protein p63.

### Type II human arrested spermatocytes display lower expression of cell cycle regulating genes

In the mouse, synapsis and DNA damage checkpoints have been shown to induce meiotic arrest analogous to human type I. Conversely, mouse models showing type II arrest have not been clearly described. However, because disturbed cell cycle regulation seems a main cause or consequence of type II arrest, we checked the current literature for mouse knockout models used in cell cycle research. Interestingly, *Ccna1*^*-/-*^ mice, which lack the gene encoding cyclin A1, display a type II-like meiotic prophase arrest without apparent problems at the pachytene stage^34, 35^. Moreover, in both mouse and human, cyclin A1 is mostly restricted to the testis and in the mouse specifically present from late pachytene till the meiotic M-phases ^36^. However, in a beeswarm plot, *CCNA1* (the human gene coding for cyclin A1) seems downregulated in both type I and II spermatocytes. Downregulation of *CCNA2* on the other hand appears more evident in type II cells (Fig. 5c). Also knockout of cyclin E2, but not cyclin E1, causes meiotic arrest in the mouse ^37^ and together both E-type cyclins control chromosome pairing, telomere stability and CDK2 localization during male meiosis in the mouse ^38^. However, in contrast the mouse, we do not find differential expression of human *CCNE2*, but instead find a clear type II specific downregulation of *CCNE1*, the gene encoding human cyclin E1 (Fig. 5c). In addition, and also specific for type II spermatocytes, we find a clear downregulation of *CDC25A* (Fig. 5c), a cell cycle regulating phosphatase whose expression has been found to be significantly decreased in a subgroup of men suffering from meiotic arrest ^39^.

### Type I and II pachytene spermatocytes fail to properly silence the sex chromosomes

Finally, we investigated whether the increased expression of *ZFY* and *ZFX* in the meiotic arrest samples could be due to disturbed meiotic sex chromosome silencing. We therefore, per included sample, plotted the number of genes expressed from the sex chromosomes, relative to the total amount of expressed genes in that specific patient sample, side by side to all included leptotene/zygotene samples (Fig. 6). Meiotic sex chromosome silencing appeared to be disturbed in both patient groups. However, the relative number of genes expressed from the Y chromosome is very limited and a two-way ANOVA analysis of the two patient groups and controls with normal spermatogenesis, followed by a Tukey’s HSD test, only demonstrated significantly disturbed meiotic silencing of the X-chromosome (adjusted p ≤ 0.0001).

**Figure 6.**
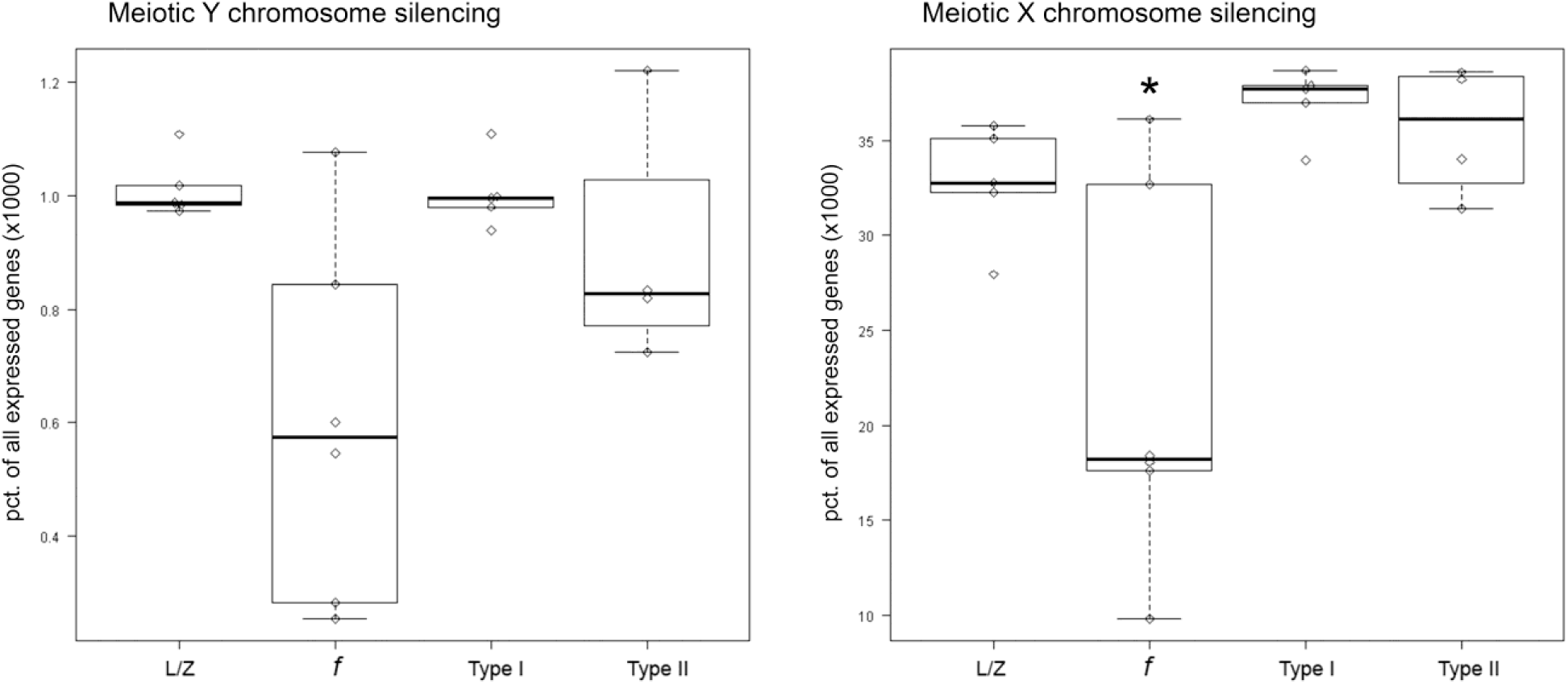
Boxplots showing the amount of genes expressed from the X and Y chromosomes relative to the total amount of genes expressed in the same sample for leptotene/zygotene (L/Z), fertile control pachytene (*f*), type I pachytene (type I) and type II pachytene (type II) spermatocytes. Significant difference between fertile and patient pachytene spermatocytes (2-way ANOVA, Tukey HSD) is labelled by an asterisk (* adjusted p ≤ 0.0001).

## Discussion

Based on microscopic evaluation of spermatogenesis and meiosis, and subsequent transcriptome analysis, we distinguish two types of meiotic prophase arrest that occur during human spermatogenesis.

Analogous to the mouse ^9, 10, 16^, human type I meiotic prophase arrest appears to be caused by incomplete synapsis of the homologous chromosomes and subsequent failure to silence the sex chromosomes; in the mouse leading to aberrant expression of the Y-chromosomal genes *Zfy1* and *Zfy2* and subsequent apoptosis of spermatocytes ^16^. Indeed, human type I spermatocytes show severe asynapsis of the homologous chromosomes, no XY-body formation and a clear upregulation of *ZFY* expression. However, we did not detect significant patient specific gene silencing of the Y-chromosome. This could be due to the fact that expression of many genes present in human pachytene meiotic cells is already established in early meiotic cells or even in spermatogonia ^32^. Because they are already expressed before chromosome silencing occurs, impaired meiotic silencing of these genes is more difficult to detect. Moreover, because the percentage of genes expressed from the Y-chromosome is below 0,1%, differences between patient groups might be too small to detect. The number of genes expressed from the X-chromosome is much higher, and indeed we observe a clear increase of X-chromosome expressed genes in both patient groups as compared to controls with normal spermatogenesis. Since the X- and Y-chromosomes are always silenced together in the meiotic sex-body, one may assume that failed X-chromosome silencing goes together with failed silencing of the Y-chromosome. Hence, analogous to the mouse synapsis checkpoint, also during human male meiosis, asynapsis of the homologous chromosomes seems to lead to increased expression of *ZFY*, most likely caused by aberrant sex-chromosome silencing.

In addition, meiotic cells can be eliminated during prophase by a separate DNA damage checkpoint including ATM, CHK2 and the DNA damage response proteins p53 or p63 ^17-20, 23^. In line with the presence of such a checkpoint, we find a clear upregulation of *TP63* (but not *TP53*) in human type I arrest spermatocytes. We did not find differential expression of putative upstream regulators of this pathway, for instance MRE11, NBS1, ATM or CHK2, in our data set. The reason for this result could be that these proteins are regulated by post-translational modifications and remain stable at the transcriptome level. Also in mouse oocytes, which do not contain a Y-chromosome or XY-silencing, the DNA damage checkpoint leads to specific activation of p63 ^20^. Moreover, it has been recently found that p53 and p63 are specifically involved in recombination-dependent mouse pachytene spermatocyte arrest ^23^. We therefore propose that human type I spermatocyte elimination can be induced by a DNA damage signaling cascade that activates p63.

While many genetic mouse models display meiotic prophase arrest analogous to the type I human meiotic arrest we describe here, only few mouse models describe a type II-like meiotic prophase arrest like we describe for type II human spermatocytes; progression further into the pachytene stage, including normal chromosome synapsis, XY-body formation and meiotic crossover formation. This is probably due to the fact that most genes studied in mouse meiosis are involved in chromosome pairing and synapsis or DSB repair, processes that when disturbed will mostly lead to type I-like arrest ^2, 3, 17^. In contrast, we selected our patient samples based on testicular histology and were thus unbiased on their genetic background. From our data it appears that the type II meiotic prophase arrest we describe for human meiosis can be defined by a decrease in transcripts required for cell cycle progression, especially the cyclins A2 and E1 and the cell cycle regulating phosphatase *CDC25A*. Indeed, in line with these data, also in mice disruption of cyclins can lead to type II-like meiotic arrest ^34, 35, 38^. The exact mechanism that underlies type II spermatocyte elimination remains to be elucidated. Considering the differential expression of genes involved in chromosome organization and RNA processing in both types of human meiotic prophase arrest, these processes could form an underlying problem for prophase meiotic arrest in general. However, in type II spermatocytes these problems may be subtler and, in contrast to type I, not directly apparent at the microscopy level. Such less apparent problems, while being overseen by the genome integrity checkpoints that induce expression of *ZFY* or *TP63* in type I spermatocytes, may still induce cell cycle arrest and subsequent type II meiotic failure. On the other hand, we have now demonstrated that, despite apparently normal chromosome synapsis and XY-body formation, also type II spermatocytes display disturbed sex chromosome silencing. However, in type II spermatocytes this seems to lead to increased expression of the X-chromosome encoded gene *ZFX*, suggesting a separate elimination pathway that is more active in type II spermatocytes.

One could argue that spermatocytes in type I and II meiotic arrest patients simply do not progress far enough to get the chance to express genes that are usually present in healthy pachytene spermatocytes. Importantly, we used the transcriptome of early pachytene spermatocytes, collected in a previous study describing gene expression throughout normal spermatogenesis ^32^, as controls. In this study, almost all genes characteristic for meiosis appeared already expressed in early spermatocytes, with early and late spermatocytes only displaying 24 DEGs. Of these 24 genes only 4, which were all downregulated in late pachytene spermatocytes in the previous study, are also found as differentially regulated in the current study. Therefore, differences in gene expression found in the arrested spermatocytes are not likely to be due to failure to reach a later pachytene stage at which these genes would normally start to be expressed.

Meiosis of one patient appeared to arrest at a metaphase stage (type III). Pachytene spermatocytes from this patient showed normal chromosome behavior and crossover formation and clustered with the controls with normal spermatogenesis after transcriptomic analysis. Considering the huge genetic and phenotypic diversity among human patients, it is not possible to delineate a list of statistically significant genes that would describe a common denominator for human male meiotic metaphase arrest from only a single patient. We therefore focused on the two types of human meiotic prophase arrest of which we analyzed the pachytene spermatocytes of 5 and 4 patients respectively.

Our study presents a comprehensive and publicly available list of genes and pathways that are involved in human meiotic prophase arrest. This list will help to understand and generate new ideas about the molecular control of human meiosis. Moreover, we identified two types of human male meiotic prophase arrest caused by distinct mechanisms. Identification and understanding of these meiotic arrest mechanisms increases our insight in how genomic stability is guarded during human germ cell development.

## Figure legends

**Extended Data Figure 1.**
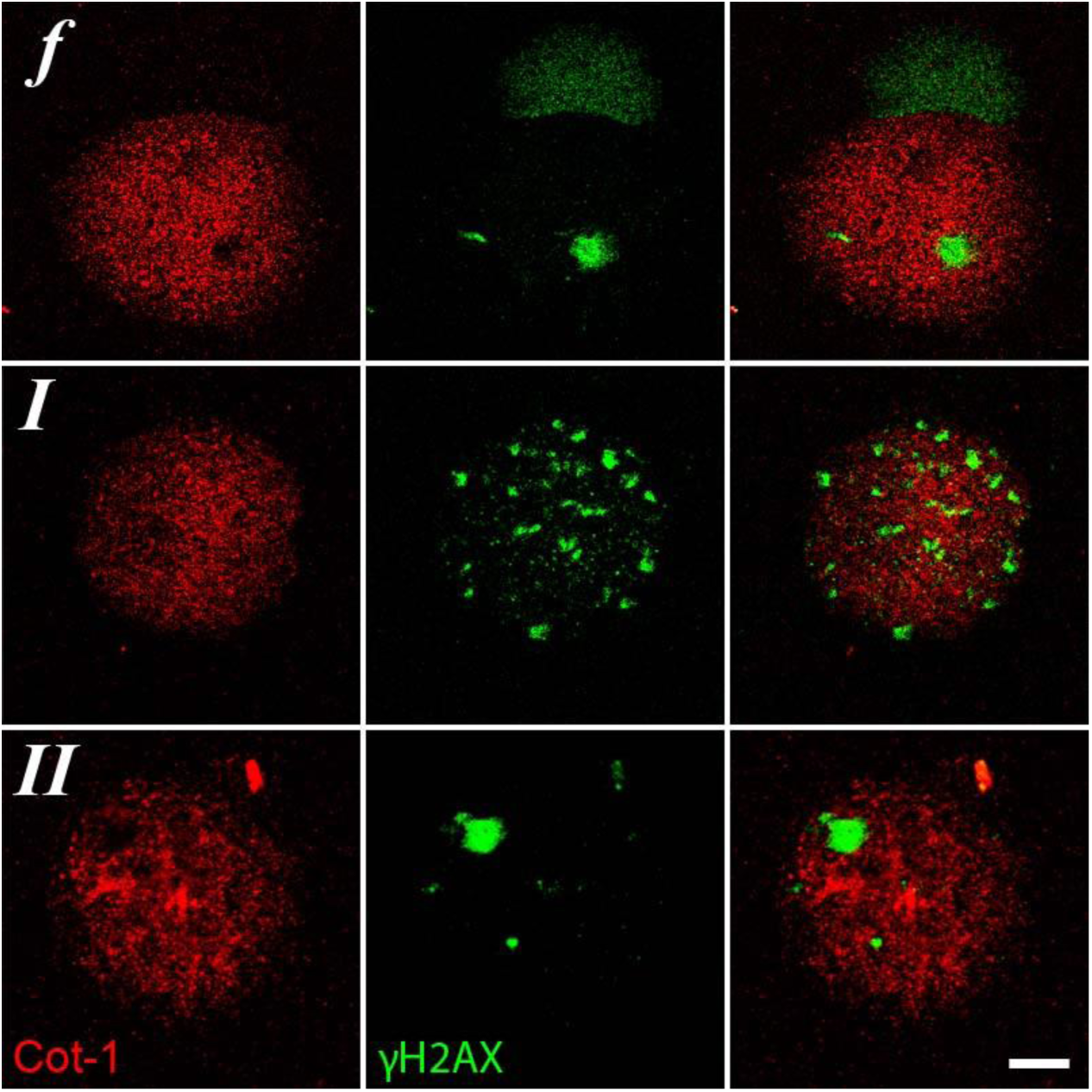
Combined RNA-FISH (Cot-1) and Immunofluorescent staining (γH2AX) of meiotic spread preparations of fertile men (***f***) and type I (***I***) and type II (***II***) meiotic arrest patients show that (i) chromosomal regions marked by γH2AX are transcriptionally silent (absence of Cot-1) and (ii) type I patients lack a transcriptionally silent XY-body (***I***). Bar = 5 μm.

**Extended Data Figure 2.**
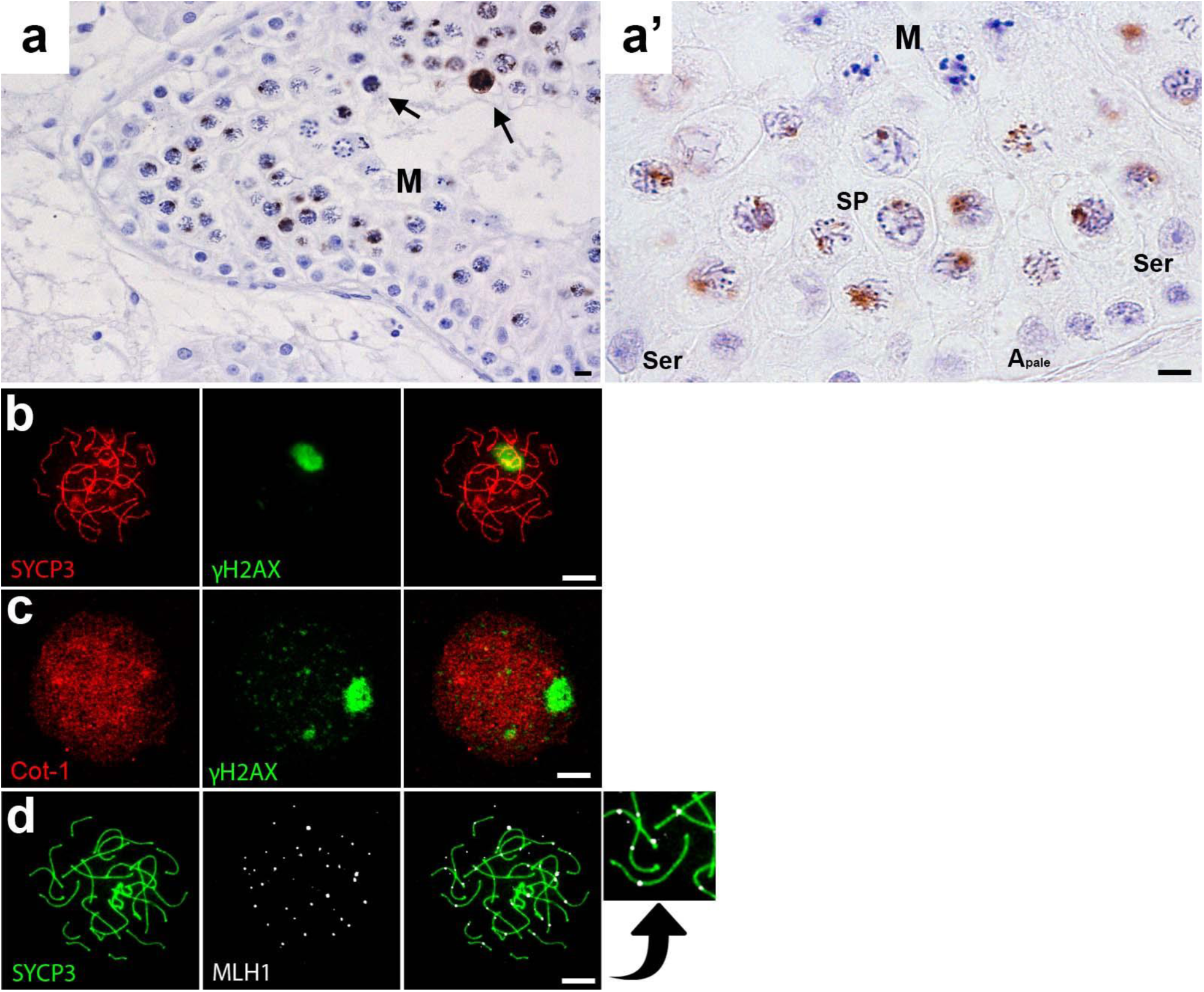
Histological evaluation and immunohistochemical localization of γH2AX in paraffin embedded testis sections of type III meiotic arrest patients (**a, a’**). These patients display normal γH2AX distribution and XY-body formation and meiotic metaphase arrest. Depicted are: Sertoli cells (Ser), A_pale_ spermatogonia, spermatocytes (SP) and meiotic metaphases (M). Bar = 10 μm. (**b**) Type III meiotic cells stained for SCYP3 and γH2AX. (**c**) Type III meiotic cells stained for Cot-1 RNA and γH2AX. (**d**) Type III meiotic cells stained for SCYP3 and γH2AX. Bar = 5 μm.

**Supplementary Table 1**. List of genes found in each k-means cluster.

**Supplementary Table 2**. Results of DAVID gene ontology analysis of genes in each k-means cluster.

## Methods

### Tissue collection

Testicular biopsies were collected with informed consent from men undergoing a testicular sperm extraction procedure (TESE) as part of their IVF treatment for subfertility at the Center for Reproductive Medicine at Amsterdam Medical Centrum (AMC). Biopsies were fixed in modified methacarn (89% methanol and 11% glacial acetic acid) and embedded in paraffin as described previously ^32^ and later used for laser capture microdissection. Remnants of the TESE procedure after sperm extraction were cryopreserved in 8% dimethyl sulfoxide (DMSO, Sigma-Aldrich) and 20% fetal calf serum (FCS) (Invitrogen) in minimum essential medium (MEM, Invitrogen) and stored at −196°C for later research use in preparing meiotic spreads. Patient IDs included in the study: AMC1805, AMC2281, AMC2489, AMC2188, URO0225, URO0229, URO0287, AMC2196, AMC2226 and AMC2062 and, for histology, AMC1849 (control patient diagnosed with obstructive azoospermia and complete spermatogenesis in all the seminiferous tubules).

### Immunochemistry

Immunohistochemical staining with mouse monoclonal anti-yH2AX (1:20,000; Millipore), on 5-μm human testis sections, was performed as described previously ^40, 41^. Human meiotic spread preparations were made according to an adapted protocol from De Vries et al. 2012 ^42^. Briefly,germ cells were isolated from testicular TESE remnants using enzymatic digestion with collagenase IV in HBBS (Hank’s Balanced Salt Solution; Gibco)/DNase solution for 20 minutes at 37°C. Subsequently, loosened tubules were incubated in a solution containing 0.25% Trypsin/EDTA diluted 1:5 in HBBS/DNase at 37 degrees till the biopsies were completely dissociated. Trypsin was inactivated using 5ml of MEM/10%FCS. Following this, the dissociated tissue was spun down at 350g for 5 minutes without break. The supernatant was removed and the resulting pellet was resuspended in SIM/TIMs. Hereafter, for spreading of the cells, the meiotic spreads protocol from De Vries et al ^42^ was followed onwards. For immunocytological staining, spreads were blocked in 1% FCS in PBS for 30 minutes at room temperature followed by an overnight incubation with the primary antibodies: mouse anti-yH2AX (1:20,000; Millipore) and goat anti-SYCP3 (1:500; R&D). The spreads were subsequently incubated for 1 hour with secondary antibodies: Alexa donkey anti-mouse 555 and Alexa donkey anti-goat 488 (Invitrogen). Slides were then counterstained with DAPI and mounted using ProLong Gold (Cell Signalling Technology). For MLH-1 staining, the combination of antibodies used was: mouse anti-MLH1 (1:500; BD Pharmingen) and goat anti-SYCP3 (1:500; R&D) and their respective secondary antibodies: Alexa donkey anti-mouse 488 and Alexa donkey anti-goat 555 (Invitrogen).

### Cot-1 DNA Fluorescence In Situ Hybridization (FISH)

To visualize nascent RNA sequences and proteins in the same sample, meiotic spread slides were first subjected to the standard immunofluorescence protocol as described above, after which they underwent a Cot-1 RNA-FISH protocol. Human Cot-1 DNA (Roche) was biotin labeled by nick translation and used as a probe, diluted in a 50% formamide hybridization mix adapted from Turner et al, 2005 ^43^ (50% formamide, 4x sodium chloride/sodium citrate solution (SSC), 20% dextran sulfate, 2mg/ml DNase/RNase-free BSA. Probe solution was denatured at 72°C for 10 minutes, chilled on ice and added to the meiotic spreads slides for an overnight incubation at 37°C. The following morning slides were washed 3x 1x SSC/50% formamide solution at 42°C followed by 3 washes with 2x SSC at 42°C and 1 rinse with 4x SSC +0.1% Tween20 at room temperature. The slides were subsequently blocked in FISH blocking solution (4x SS, 0.1%Tween 20, 4mg/m DNase/RNase-free BSA for 30 minutes at 37°C. Slides were then incubated with avidin–Cy3 (1:5000, Jackson ImmunoResearch). To enhance the signal slides were then incubated with biotinylated anti-avidin (1:500, Vector Lab) followed by a final incubation with avidin-Cy3, each incubation was for 30 minutes at 37°C. Finally, the slides were stained for yH2AX and SCP3 as described above.

### Microscopy

Bright field microscopy images were acquired at room temperature using an Olympus BX41 microscope equipped with an Olympus DP20 color camera. Fluorescence microscopy images were acquired at room temperature using a Plan Fluotar 100x/1.30 oil objective on a Leica DM5000B microscope equipped with a Leica DFC365 FX CCD-camera. Images were analysed using Leica Application Suite Advanced Fluorescence (LAS AF) software. The presented figures were constructed using Adobe Photoshop CS5 version 12.0.

### Single cell laser capture microdissection and RNA preparation

Directly prior to laser dissection microscopy (LDM), 5μM thin sections of testis tissue were mounted on Superfrost glass microscope slides (Thermo Scientific) and HE-stained as described previously ^32^. 500 pachytene spermatocytes per patient were individually laser dissected and pooled as described previously ^32^. Laser dissected cells were captured in silicon coated adhesive caps (Adhesive cap 500 opaque tube, Zeiss) and were lysed at 42°C in 10μl of extraction buffer provided in the PicoPure RNA isolation kit (Arcturus). Cell lysates were stored at −80°C until further use. All procedures were done under RNase free conditions. Total RNA was isolated from cell lysates using the PicoPure RNA isolation kit (Arcturus) according to the manufacturers’ protocol including an on-column DNase treatment. RNA was eluted in 10μl elution buffer. Subsequently, the RNA was concentrated to a volume of 5 μl with a speed vacuum centrifuge for 8 minutes. Total RNA isolated from 500 pooled cells was SPIA-amplified using the Ovation RNAseq V2 System (Nugen) as described previously ^32^.

### RNA sequencing

The amplified cDNA was sheared using the Covaris S220 (Thermo Fisher Scientific). DNA libraries were made from the SPIA-amplified cDNA and sequenced single-end, 50bps on the HiSeq2000 Illumina platform obtaining at least 10 million reads using 8 pmol per library.

### Bioinformatics

Bioinformatic analysis was carried out using a pipeline previously ^32^. Briefly, samples with a normalization factor between 0.6-1.4 were included in the analyses. For multidimensional scaling analysis, comparing previously derived germ cell transcriptomes with arrested pachytenes ^32^, all samples included in the plot were normalized together. For all further analyses, the fertile leptotene/zygotene, pachytene, type I and II spermatocytes were normalized together. A gene was considered expressed if it had >1 count per million present in at least three individuals per sample group. Raw counts were transformed to moderated log-counts-per-million before filtering using the cpm function with default parameters. A list with differentially expressed genes (DEGs) between the spermatocytes was obtained by estimating the mean-variance of the log counts using the *voom* method and analysing these with the empirical Bayes pipeline as implemented in *limma* (version 3.22.7). After correcting for multiple testing, a p-value of <0.05 was considered significant for DEG analysis. K-means clustering (default algorithm) was used to obtain plots for the scaled normalized relative gene expression data on a log scale for each expressed gene using packages *clValid* (version 0.6-6), *cluster* (version 2.0.2) and *stats*. Gene ontology analysis was performed using the functional annotation clustering tool in DAVID. An enrichment score of >1.3 was considered significant.

#### Acknowledgements

This work was supported by ZonMw VIDI-grant 91796362 to S.R. and an AMC Fellowship and The People Programme (Marie Curie Actions) of the European Union’s Seventh Framework Programme (CIG 293765) to G.H.. We thank Dr. R. Kerkhoven of the Genomics Core Facility of the Netherlands Cancer Institute, Amsterdam for conducting the RNA sequencing. We thank Marieke de Vries for demonstrating her protocol for meiotic spreads from frozen human testis biopsies. We thank Daisy Picavet and the core facility Cellular Imaging of the AMC for assistance with confocal microscopy and use of their equipment. We thank Andreas Meissner (Department of Urology, Academic Medical Center, Amsterdam, The Netherlands) for providing us with patient material.

## Author Contributions

S.Z.J., A.M.M.P., S.R. and G.H. conceived and designed the study. S.Z.J., C.M.K. and S.K.M.D. conducted all experiments. S.Z.J. and A.J. performed bioinformatic analyses. S.Z.J., A.J. and G.H. interpreted the results. S.Z.J., A.J. and G.H. performed data visualization. S.Z.J. and G.H. wrote the manuscript. S.Z.J., A.J., S.R., A.M.M.P. and G.H. critically read the manuscript.

## Author Information

All sequence data have been submitted to NCBI (SRA) and are available under the accession number: PRJNA373978. The authors declare no competing financial interests. All correspondence should be addressed to Geert Hamer.

